# Effects of gabapentin on ongoing behaviors displayed by mice with chemotherapy neuropathy

**DOI:** 10.64898/2026.06.24.734356

**Authors:** Bradey A. R. Stuart, Bhavya S. Dharanikota, Cheryl L. Stucky

## Abstract

Chemotherapy-induced peripheral neuropathy (CIPN) is a common and painful side effect of paclitaxel (PTX) treatment. The most common measures of painful neuropathy focus on evoked mechanical hypersensitivity, but clinically relevant ongoing pain remains understudied in preclinical models. Automated machine learning methods for pose estimation and behavioral classification have been proposed to capture non-evoked pain-like behaviors, though these approaches have primarily been applied to unilateral injury models such as spared nerve injury or unilateral inflammatory compound injection. Here, we evaluated the extent to which paclitaxel-induced CIPN affects the posture and spontaneous behavior of freely moving mice using a commercially available automated recording system (BlackBox). We found that paclitaxel-treated mice develop a broad and reproducible behavioral and postural phenotype relative to vehicle-treated controls, characterized by reduced front paw luminance and print size, increased front paw lifting, and altered body measurements consistent with a guarded posture. This phenotype was replicated across two independent cohorts and was detectable at both day 2 and day 6 following the final paclitaxel injection. To identify behavioral features specific to CIPN, we administered gabapentin, an analgesic often used to treat neuropathic pain in patients, to determine whether paclitaxel-induced behavioral changes could be attenuated. Gabapentin reduced several behavioral features in both paclitaxel-treated and vehicle-treated animals, suggesting that its effects on posture and gait are not specific pain in CIPN. These findings demonstrate that automated behavioral recording captures a robust paclitaxel-induced postural phenotype but question whether captured behaviors are indicative of ongoing pain as alleviated by gabapentin.

## Introduction

In animal models, pain has commonly been studied using experimenter-evoked behavioral responses to normally innocuous or noxious stimuli[30]. While these evoked measures have been invaluable for understanding peripheral and central sensitization, their clinical translation has been limited, and patients with chronic pain conditions frequently report non-evoked spontaneous or ongoing pain as their primary concern[20,30]. Pre-clinically, this dimension of the pain experience remains understudied. Historically, non-evoked pain-like behaviors have been assessed using grimace scores[8,22], burrowing behavior[14,37], gait analysis[10,35], weight bearing asymmetry[19,27], and conditioned place aversion[17,33]. Automated scoring of these methods has substantially increased the throughput and objectivity of several of these techniques. Recent advances in machine learning have enabled automated pose estimation and behavioral classification in freely behaving animals, opening the possibility of unbiased characterization of pain-related behaviors[38]. Unilateral injury models such as spared nerve injury and inflammatory pain have been particularly well studied, where asymmetric postural changes provide a clear readout of pain-related compensation[38]. However, their application to systemic pain conditions such as chemotherapy-induced peripheral neuropathy (CIPN), where postural changes may be more subtle, remains limited.

Paclitaxel (PTX) is among the most widely used chemotherapeutic agents for the treatment of solid tumors, but its clinical utility is frequently limited by the development of CIPN, a painful and often dose-limiting side effect that affects up to 70% of treated patients[3,9]. Paclitaxel hyper-stabilizes microtubules, leading to axonal dysfunction and damage in both large and small diameter sensory neurons[5,31], ultimately producing a painful peripheral neuropathy characterized by tingling, numbness, and spontaneous pain in the distal extremities. In rodent models, paclitaxel reliably induces mechanical hypersensitivity and cold allodynia[21], but whether it produces detectable changes in non-evoked spontaneous behavior and posture has not been comprehensively characterized using automated behavioral recording approaches.

Here we evaluate the hypothesis that paclitaxel-induced CIPN produces a measurable and reproducible postural and behavioral phenotype in freely moving mice. We further hypothesize that administration of gabapentin, an analgesic used clinically for neuropathic pain, will selectively reduce pain-associated behavioral features, providing a pharmacological tool to distinguish pain-specific postural changes from general treatment effects. Together these experiments aim to validate automated behavioral recording as a tool for characterizing the non-evoked behavioral consequences of CIPN.

## Results

To investigate the role of CIPN on non-evoked behavior in mice we utilized a recently developed technique to evaluate the posture and behavior in freely behaving mice, commercially termed BlackBox. In this behavioral device, a camera situated beneath mice captures transmitted light to capture posture. Additionally, the device utilizes infrared red light to measure paw luminance, which is correlated with paw pressure and weight bearing[38]. Following a 4-dose induction of CIPN with paclitaxel (PTX), we performed behavioral recordings on day 2 and day 6 following the last injection (Fig 1A). This PTX dosing scheme reliably induces robust evoked pain like behaviors beginning after the first injection, that persists for multiple weeks following cessation[12,23]. Day 2 and day 6 were chosen to capture behavioral phenotypes outside of acute administration related changes. To confirm mice experience an ongoing pain during this time, we performed automated facial grimace analysis[22], which has been shown to be elevated during ongoing pain, and found an increase in the mean grimace score, as well as the number of frames with a high (>5 facial action units) grimace score (Supplemental Fig 1A,B). These results suggest that PTX treated mice experience ongoing pain during the chosen recording time frame. During Blackbox recordings, the output after automatic analysis consists of 65 features that could be classified as body angles, body measurements, luminance, paw lifting, pressure indices, or paw print size. Body measurements and angles are calculated from automatic pose estimation (Fig 1B), while luminescence, pressure indices, and print size are calculated using infrared light measurements (Fig 1B). To better understand the overall behavioral differences between CIPN and vehicle animals we performed a permutation-based multivariate analysis of variance (PERMANOVA) using postural and behavioral features collected at day 2 and day 6 following the last injection of paclitaxel. PERMANOVA revealed a significant effect of day (F[1,38] = 1.86, R^2^ = 0.047, p = 0.032), indicating that the overall behavioral profile changed across timepoints regardless of treatment. When modeling day and treatment together, a significant proportion of behavioral variance was explained (F[2,37] = 3.87, R^2^ = 0.173, p = 0.029), suggesting that treatment group differences contribute meaningfully to the multivariate behavioral phenotype beyond the effect of time alone. The day × treatment interaction trended toward but did not reach significance (F[3,36] = 2.84, R^2^ = 0.191, p = 0.060), suggesting a possible progressive change in behavioral phenotype between paclitaxel and vehicle groups over time, although this was not statistically significant.

**Figure 1:**
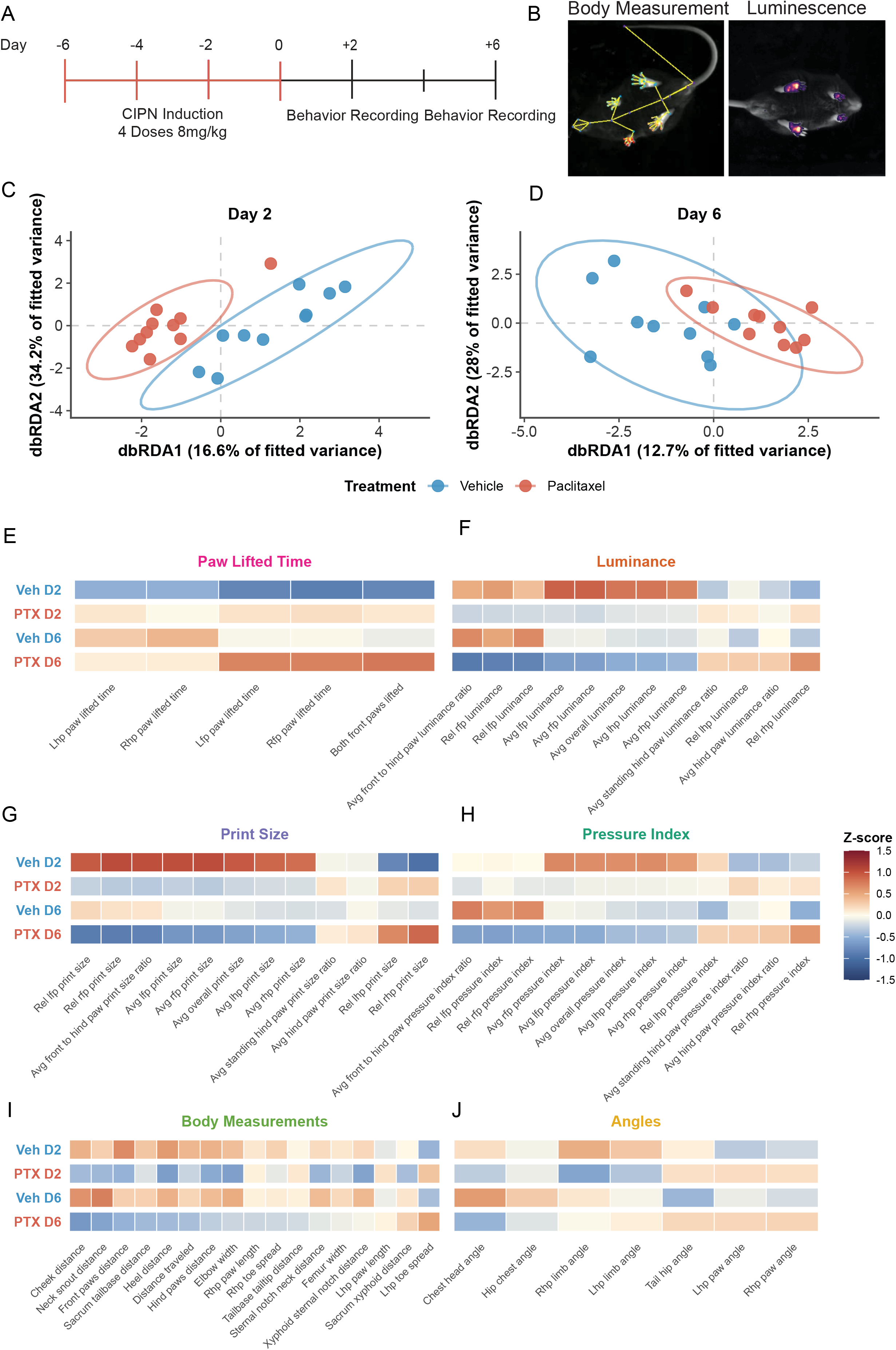
Automated scoring of paclitaxel induced CIPN reveals a distinct postural and behavioral phenotype. A) Experimental timeline for induction of CIPN and behavioral recording days. B) Example of how body measurements are calculated (left) and paw pressure (luminescence, right) measures look after processing raw videos. C, D) Dimensionality reduction using Distance-based Redundancy Analysis (dbRDA) on day 2 (C) and day 6 (D), reveals distinct separation of PTX and vehicle treated animals. Statistical analysis (PERMANOVA) revealed a significant revealed a significant effect of day (F[1,38] = 1.86, R^2^ = 0.047, p = 0.032). Day and Treatment (F[2,37] = 3.87, R^2^ = 0.173, p = 0.029. Qualitative heatmap analysis of Paw lifted time (E), luminance measures (F), print sizes (G), Pressure indices (H), Body measurements (I) and body angles (J).

To visualize these results, redundancy analysis (RDA) constrained by day and treatment was performed, revealing distinct separation of paclitaxel-treated animals from vehicle controls at day 2 (Fig 1C) and modest separation at day 6 (Fig 1D), consistent with the statistically significant PERMANOVA results (Fig 1A). To better visualize the entire 65 feature dataset, we averaged each group’s mean, calculated a z-score, and displayed these results as a heatmaps for each category of behavior (Fig 1B). Variables within each category were sorted by mean z-score to highlight the most differentially abundant features. Paclitaxel-treated animals showed a broad pattern of elevated z-scores relative to vehicle controls across multiple feature categories, with differences on both days 2 and 6. Of note, the paw lifted time category showed consistent elevation in paclitaxel animals at both day 2 and day 6 in the front paw lifting behavior (Fig 1E). For the category of luminance, paclitaxel treated mice displayed decreases in front paw luminance and decreases in the front to hind paw luminance ratio (Fig 1F). Furthermore, although the PTX treated animals displayed raw luminance value decreases in the individual left and right hind paws (Fig 1F), the PTX treated mice displayed increased luminance while standing on their hind paws (Fig 1F). These results are consistent with decreased contact of paws on the front paw (ie. paw lifting, Fig 1E) with a corresponding increase in weight shifted to their hind paws. Looking at the total size of the paws, there was an overall decrease in the measured front paw size (Fig 1G) and an overall increase in hind paw print sizes (Fig 1G) compared to vehicle animals on day 2 and day 6. For the pressure index category, which represents the raw luminance value divided by the print size, showed broad reductions (Fig 1H). Specifically, the front paw pressure index was decreased on day 2 and day as well as the overall pressure index of all four paw (Fig 1H). Together this suggests not only a weight shift from front paws to hind paws, but that in general the PTX treated animals tended to place less pressure on each paw specifically. For the category of body measurements (Fig 1I) PTX treated animals showed modest decreases across many measurements, including decreases in cheek distance, neck to snout distance, heel distance, and front paw distance (Fig 1I). Variables such as toe spread were not consistently affected by PTX compared to vehicle. For the body angle measurements, measured variables were mostly unaffected (Fig 1J). Despite decreases in hind paw limb angles, this was differently affected on day 2 vs day 6 in PTX treated animals (Fig 1J), with overall small effect sizes. Overall, this qualitative analysis of the features across categories through heatmaps shows a coordinated shift in postural and gait features in paclitaxel-treated animals, with variables suggestive of a shift to increased hind paw weight and increased front paw lifting behavior.

To specifically identify features that differed between CIPN and vehicle-treated animals, linear mixed effects models were applied to each of the 65 behavioral features independently, with day, treatment, and their interaction as fixed effects and mouse as a random intercept. Following Benjamini-Hochberg correction to control the false discovery rate (FDR < 0.05), 20 of 65 behavioral features survived multiple comparisons. Significant features were driven by a main effect of treatment rather than a day × treatment interaction, indicating that paclitaxel-induced behavioral changes were stable across both timepoints rather than progressive. To visualize the direction and magnitude of treatment effects across all 65 features, effect sizes (Cohen’s d) were plotted against FDR-corrected p-values, revealing that significant features (-log_10_ padj >1.3) spanned multiple behavioral categories and were characterized by both increases (Cohen’s d > 0) and decreases (Cohen’s d < 0) relative to vehicle controls (Fig 2A). Overall, we found 20/65 variables that survived multiple comparisons and were highly statistically significant. Consistent with our qualitative observations (Fig 1E-J), we observed highly statistically significant increases across the paw lifted time category (Fig 2B-D). Additionally, across the luminance values, we observed consistent decreases on days 2 and 6 (Fig 2E-I). For print sizes, again we observed decreases across the overall print size, and the front paws (Fig 2J-N) and corresponding increases in the ratio of the hind paws (Fig 2O,P). Variables in the body measurements category that survived multiple comparisons included decreases in heel distance, front paw distance, cheek distance, and the neck to snout distance (Fig 2R-U). All together the combination of increased paw lifting, decreased paw pressure (luminance) with decreases in front paw print size is consistent with a putative guarded position, in which mice shift weight from their front to hind paws, and stand predominantly on their hind paws. Treatment effects were consistent across day 2 and day 6 for all significant features, in line with the absence of a significant day × treatment interaction. Clinically, because PTX is often used to treat ovarian and breast cancers, PTX is disproportionately used more often in women[11]. Therefore, we compared the magnitude of paclitaxel’s effect size on each feature in male and female mice, and found that in general, behavioral features that are increased in males are increased in females (Pearson correlation coefficient R = 0.65, p<0.001) (Supplemental Figure 1C). As such, we chose to analyze these findings aggregated. Together, these findings indicate that paclitaxel treatment produced a broad, body wide behavioral and postural phenotype characterized by front paw lifting, reduced front paw contact, and altered body measurements that were detectable at both day 2 and day 6 following the final injection, in both males and females.

**Figure 2:**
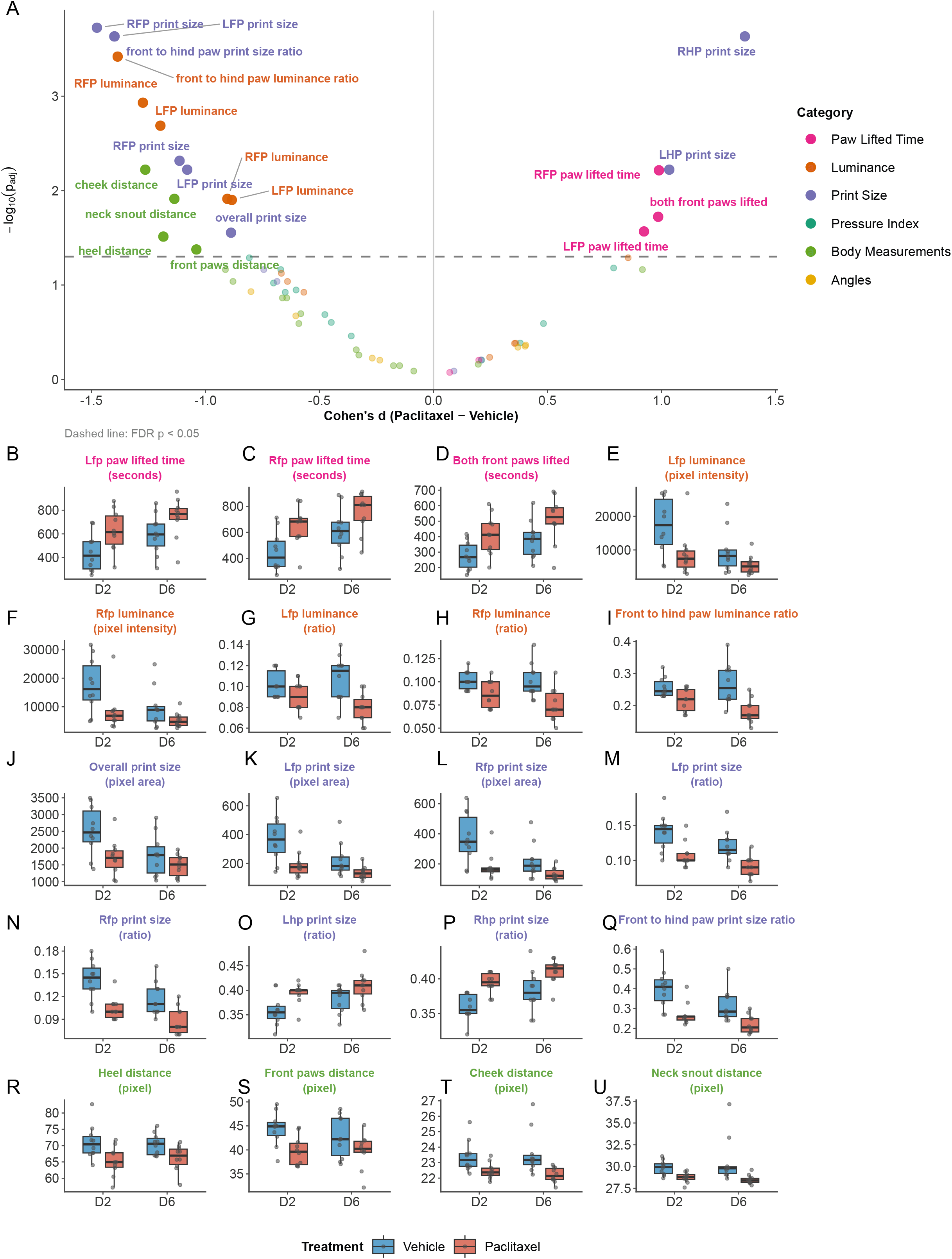
Univariate analysis of automated behavior analysis identifies 20 variables increased or decreased in CIPN animals. A) Volcano plot of all variables using Cohen’s D (average of paclitaxel – vehicle) animals and log transformed benjamini hochberg corrected p-values with FDR <0.05 reveals highly statistically significant variables in paw lifted time, luminance, print size, and body measurement categories. (B-U) Time course visualization of each of the 20 identified variables surviving multiple comparisons (A). n=20 mice (10 PTX (5M/5F), 10 vehicle (5M/5F)

To determine which possible behavioral or postural changes are putative pain like behaviors, we hypothesized that administration of a known analgesic would attenuate the behaviors with CIPN. Although there are no current FDA approved treatments for CIPN pain, gabapentin has shown efficacy in treating pain symptoms in some patients with CIPN[25]. Additionally, gabapentin has been shown to reduce spontaneous pain behaviors in sickle cell disease mouse models[29], and mechanical hypersensitivity in other pain models[18]. Thus, we chose gabapentin as an analgesic to evaluate changes in behavior following gabapentin administration. We induced CIPN in a separate cohort of 20 mice, and on days 2 and 6 we administered gabapentin or vehicle intraperitoneally, recording both pre and post analgesic behaviors. We conducted recordings 45 mins after administrations, to capture the 1-hour post-administration timepoint, which corresponds to the time with the highest serum concentration of gabapentin in mice[16]. First, we asked whether the significant variables identified in Figure 2 were recapitulated in this cohort of mice.

To assess whether the paclitaxel-induced behavioral phenotype was reproducible across independent cohorts, effect sizes were calculated (Cohen’s D, Fig 2A) and compared between the initial experiment and the second cohort, prior to gabapentin administration. The PTX-induced behavioral phenotype was highly reproducible across both experiments on both days. At day 2, effect sizes of the previously identified 20 significant variables, were strongly correlated between experiments (Pearson r = 0.954, p < 0.001) (Fig 3A), indicating that the direction and relative magnitude of paclitaxel-induced changes were consistent across cohorts. At day 6, the correlation remained strong (r = 0.877, p < 0.001) (Fig 3B), confirming that the phenotype was maintained at the later timepoint in both experiments. Features showing the most consistent replication across both timepoints included paw lifted time measures, and luminance and print size ratio features, which showed large negative effect sizes in both experiments (Fig 3A,B). Of note, most features tended to have lower effect sizes in the second cohort, suggesting that despite high correlation, the degree to which each feature is increased or decreased in CIPN animals may vary between experimental conditions. Overall, these findings highlight that paclitaxel reliably induces a consistent non-evoked behavioral phenotype across independent animal cohorts.

**Figure 3:**
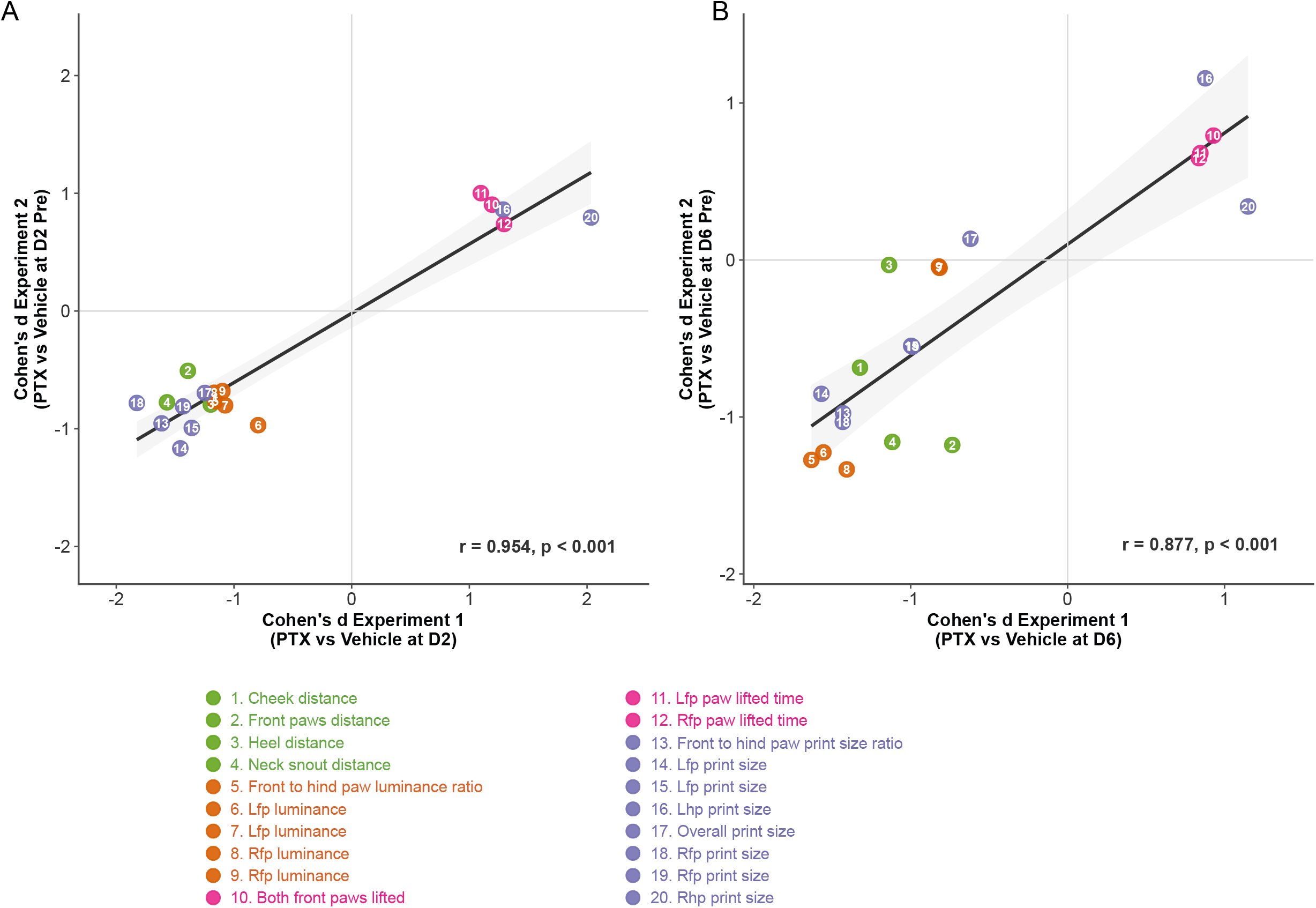
Correlation analysis of identified variables reveals high level of reproducibility between cohorts. Using a second cohort of animals, a comparison of eUect sizes (Cohen’s D PTX-Veh) on Day 2 (left), and day 6 (right) highlights a high correlation between identified variables in Figure 2.

Visualizing the 20 significant features on heatmaps separated by category and displaying mean values as z-scores relative to each group’s own baseline (Fig 4A-D) confirmed that paclitaxel-treated animals showed broad and consistent deviations from baseline across paw lifting, luminance, print size, and body measurement categories at both day 2 and day 6 pre-gabapentin timepoints. Notably however, gabapentin produced significant changes in more features in vehicle animals (15/20) than in paclitaxel animals (8/20) (Fig 4A-D), suggesting that gabapentin broadly affects posture and gait in a manner that is not specific to the paclitaxel-induced phenotype.

**Figure 4:**
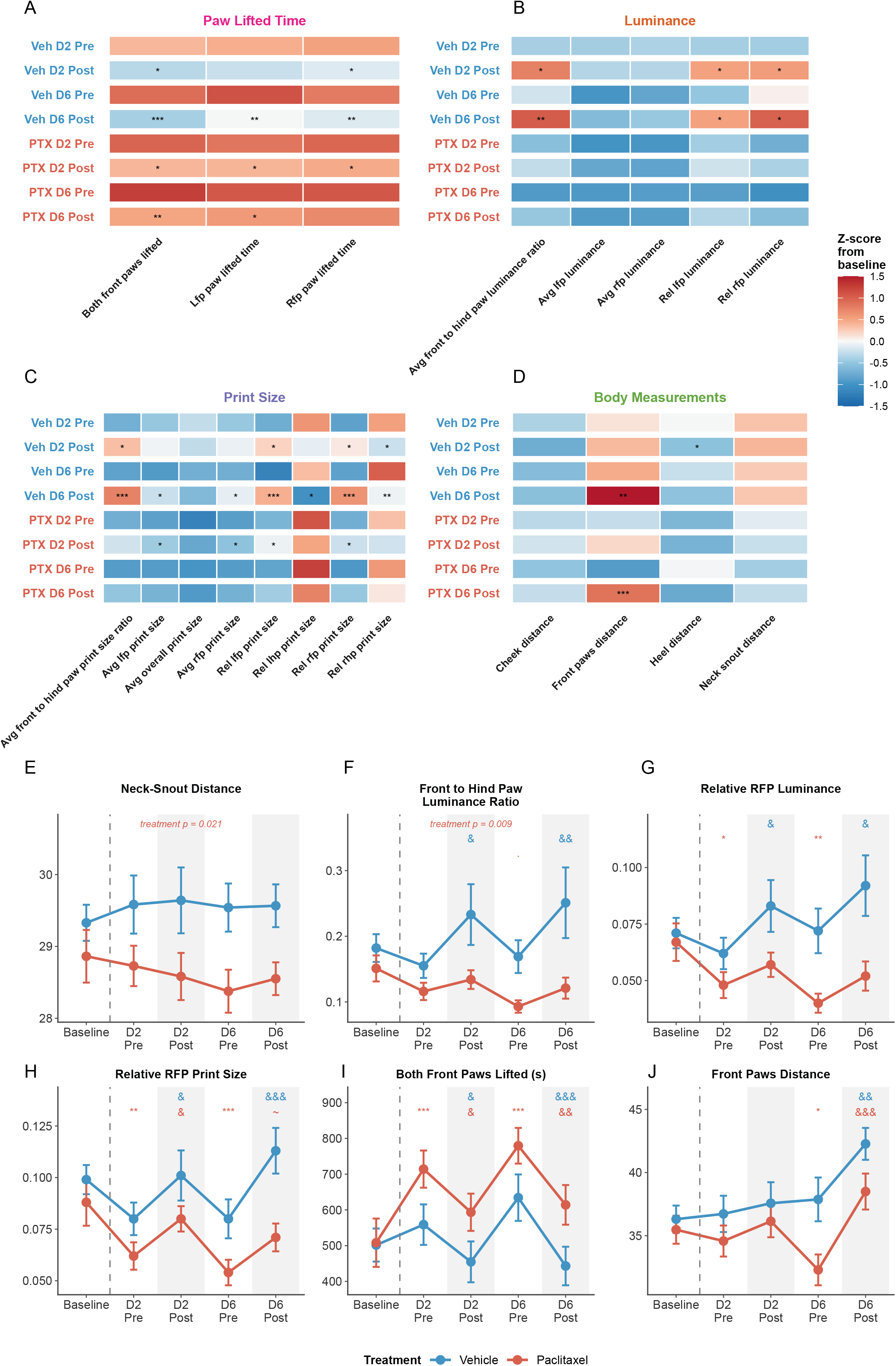
Exposure to gabapentin produces varied responses in naïve and CIPN animals. Following treatment with gabapentin, heatmap of z-scores for 20 variables identified in Fig 2 relative to each groups baseline, separated by paw lifed time (A), luminance (B), Print Size (C) and body measurements (D) (* <0.05,**<0.01,***0.001, & <0.05 for gabapentin vs pre-exposure) (E-J) Trajectory plots of six representative behavioral features across all five timepoints: baseline, day 2 pre-gabapentin (D2 Pre), day 2 post-gabapentin (D2 Post), day 6 pre-gabapentin (D6 Pre), and day 6 post-gabapentin (D6 Post). Features shown are neck-snout distance (E), front-to-hind paw luminance ratio (F), relative RFP luminance (G), relative RFP print size (H), both front paws lifted time (I), and front paws distance (J) (* <0.05,**<0.01,***0.001 for comparison to baseline, & <0.05, &&<0.01, &&&<0.001, ~ p<0.1 for Gabapentin contrast relative to pre-exposure). n=20 mice (10 PTX (5M/5F), 10 vehicle (5M/5F)

For some features gabapentin produced no detectable changes in either PTX or vehicle animals. For example, neck-snout distance was reduced in paclitaxel animals across the experiment (treatment main effect p = 0.021, Fig 4E) though individual timepoint contrasts did not reach significance, and gabapentin produced no significant changes in this feature in either group (Fig 4D, E).

For other features gabapentin uniquely produced changes only in vehicle animals. The front-to-hind paw luminance ratio increased following gabapentin in vehicle mice at both day 2 post (p < 0.05) and day 6 post (p < 0.01, Fig 4B, F), reflecting a shift toward greater front paw pressure, while paclitaxel mice showed no significant gabapentin response at either timepoint. Relative RFP luminance was significantly reduced in paclitaxel animals at day 2 pre-gabapentin (p < 0.05) and day 6 pre-gabapentin (p < 0.01) relative to baseline with no significant gabapentin effect, while vehicle mice showed significant increases following gabapentin at both day 2 post (p < 0.05) and day 6 post (p < 0.05, Fig 4B, G). Similarly, gabapentin uniquely modified the front-to-hind paw print size ratio (Fig 4B), relative left and right hind paw print sizes (Fig 4C), and heel distance (Fig 4D) in vehicle animals only.

For some features, however, gabapentin produced effects in both vehicle and paclitaxel animals. Relative RFP print size was significantly reduced in paclitaxel animals at day 2 (p < 0.01) and day 6 pre-gabapentin (p < 0.001) relative to baseline (Fig 4C, H). Gabapentin produced a significant increase in print size in vehicle mice at day 6 post (p < 0.001) and a non-significant trend in paclitaxel mice (p < 0.1), suggesting that gabapentin effects on print size are more pronounced in vehicle than paclitaxel animals. Both front paws lifted time was significantly elevated in PTX animals at day 2 pre-gabapentin (p < 0.001) and day 6 pre-gabapentin (p < 0.001), as observed previously (Fig 2A, B; Fig 4A, I). Gabapentin reduced paw lifting in paclitaxel mice at both day 2 (p < 0.05) and day 6 (p < 0.01) but also reduced paw lifting in vehicle mice at day 2 (p < 0.05) and day 6 (p < 0.001), indicating that this reflects a general gabapentin effect on paw lifting rather than reversal specific to the PTX phenotype.

Front paws distance provides a notable exception that may reflect a partially CIPN-specific gabapentin response. Paclitaxel animals showed significantly reduced front paws distance at day 6 pre-gabapentin relative to baseline (p < 0.05, Fig 4D, J), while vehicle animals showed no significant deviation from baseline prior to gabapentin. Gabapentin administration significantly increased front paws distance in paclitaxel animals at day 6 post (p < 0.001) to baseline levels. While gabapentin also increased front paws distance in vehicle mice at day 6 post (p < 0.01), the combination of a pre-existing deficit and its complete reversal by gabapentin is unique to the paclitaxel group.

Together these findings demonstrate that gabapentin produces postural effects that are largely not specific to the paclitaxel-induced behavioral phenotype, with most gabapentin-sensitive features similarly affected in vehicle and paclitaxel animals. The absence of consistent phenotype-specific reversal suggests that the behavioral features captured by automated recordings suggest that gabapentin at this dose and timepoint, suggests that the altered behaviors may not be pain specific changes in CIPN.

## Discussion

Here we find that PTX induced CIPN produces a consistent and repeatable non-evoked phenotype when animals are monitored in the dark and in the absence of an observer. Notably, we recorded animal behaviors up to 6 days following the last injection of PTX, a time point in which PTX is largely gone from mouse serum[24]. Although it is known that mechanical hypersensitivity in CIPN animals persists for many weeks, it is completely unknown what non-evoked behaviors persist after chemotherapeutic exposure. In this study we specifically found an overall CIPN phenotype that indicates a hunched or guarded position, consisting of weight bearing shifts to the hind paws, an increase in front paw lifting behavior that is accompanied by a decrease in front paw distance. We interpret this decrease in front paw distance as potentially indicative of grooming behavior or possibly another behavioral shift that includes the paws close together. The use of unbiased classifications of key points or author automated behavior classification may aid in the identification of the specific behavior altered in CIPN[13,34]. Additionally, this study is unique in using automated posture and behavioral identification in a non-unilateral injury model. Prior work has used models of pain from formalin, ultraviolet burn, and capsaicin injections[4,38]. To our knowledge, only one study has used a potentially non-unilateral injury model of spinal cord injury[15]. However, this mechanical trauma model still produces focal deficits below the lesion, in contrast to CIPN, a systemic model for painful neuropathies. Another benefit our study is that uses relatively long (25 minute) recordings, which to our knowledge have not been utilized before, but offer the benefit of capturing what may be rare behaviors.

One advancement of this study is that we take a step toward advancing pain models to better recapitulate patient phenotypes. Notably, treatments for preclinical models of pain including CIPN have largely failed in clinic due to the inherent differences in how we measure pain[7,32]. Historically, there has been a rampant increase in the use of mechanical hypersensitivity tests as a proxy for pain like behaviors[30], particularly in the hind paw. However, patients with CIPN report pain in burning in the hands and feet. Anatomically, the front paws have been neglected by pain researchers. As such, the finding in this study that CIPN mice shift to their hind paws, may suggest aversion of front paw touch to a surface. This is quite similar to patient mechanical hypersensitivity, who do not get touched by von Frey hairs, but report pain when putting on clothes. Although non-evoked pain like behaviors exist, they are still relatively uncommonly used, due to technical or throughput limitations. Our findings also show that consistent with previous literature, the acute exposure to the analgesic gabapentin at a clinically relevant dose (30mg/kg) is sufficient to induce behavioral changes in mice, in both CIPN or vehicle animals. While this study did not find a specific behavior unique to CIPN animals that was attenuated by the analgesic gabapentin, the front paws distance attenuation raises the possibility that some features may capture gabapentin-sensitive pain-related postural changes. It is possible that the mechanism by which gabapentin produces analgesia is through common pathways to naïve or injured animals. It is also possible that the observed effects may be due to gabapentin induced movement side effects, namely sedation which are often seen clinically[26,36], however, this is less likely at the does chosen for this study as has been shown previously[1,28,29]. Additionally, this study is limited by the acute exposure to the analgesic gabapentin. Patients taking gabapentin for neuropathic pain take daily medication. This study emphasizes the need for additional pharmacological validation of analgesics with distinct mechanisms, across different timescales and dosing treatments, to begin to decipher the complexity of ongoing pain behaviors.

## Materials and Methods

### Animals

All animals were 8 week old C57/Bl6 animals purchased from JAX(Strain #:000664) Equal numbers of male and female mice were used for all experiments (5M/5F per group/treatment). All mice were group housed by sex, maintained on a 12:12 light/dark cycle and bred in a climate-controlled room, with *ad libitum* access to food and water. All protocols were conducted in accordance with the National Institutes of Health guidelines and were approved by the Institutional Animal Care and Use Committee at the Medical College of Wisconsin (Milwaukee, WI; protocol 383).

### Induction of CIPN

CIPN was induced as we and others have done previously[2,6,23]. Paclitaxel (Taxol, European Pharmacopoeia, Sigma-Aldrich) was dissolved in vehicle (16% ethanol, 16% Cremophor, and 68% saline) to achieve a working concentration of 0.8mg/ml. Mice received intraperitoneal injections of paclitaxel (8mg/kg) or vehicle every other day for a total of 4 injections.

### Animal Behavior

#### Blackbox Behavior Recordings

Animals were allowed one week to acclimate following arrival to the MCW animal facility. After 1 week, animals were introduced to behavioral room and allowed to acclimate for 1-2 hours. Following acclimation to the room, animals were acclimated to behavioral apparatus for 30 minutes. For recordings, animals were placed inside BlackBox behavioral chamber for 25 mins. During the time in which recording metadata were entered into the computer (approximately 2-3 minutes), the experimenter observed if any mouse had urinated in its recording chamber, which has been shown to change luminance measurements[38]. If an animal did it was briefly removed and the surface was wiped dry. Habituation and this observation period dramatically reduced the number of animals which urinate during recording. The same experimenter conducted all experiments to minimize inter-experimenter induced phenotypes. Experimenter was not present in room during behavior recordings of non-evoked behavior. As behavior is automatically processed, captured, and analyzed independent of observer, experimenter was not blinded to animal conditions.

#### Automated Facial Grimace recordings and analysis

Mice were placed in 3-D printed behavior boxes with an open front (5” x 3” x 3”) for 30 minutes to acclimate. 30 minute recordings were captured using Sony CX405 video camera. Recordings were conducted without the experimenter in the room. Grimace videos were uploaded to the Painface© server for automated scoring, which calculates the mean grimace score and percentage of frames with a grimace score >5 (high grimace state)[22].

### Dimensionality reduction for visualization of group differences

To visualize the multivariate behavioral structure constrained by the experimental factors of interest, redundancy analysis (RDA) was performed using the rda function in the vegan package (vegan : 2.7.3) in R (4.5.2). The analysis was conducted on a standardized behavioral matrix comprising 65 measures of gait, posture, and locomotion, with standardization performed by z-scoring each variable across all individual animal observations prior to analysis. The RDA axes were constrained by day and treatment group, such that the resulting ordination reflects only the proportion of behavioral variance attributable to these experimental factors. Site scores for each animal at each timepoint were extracted and plotted, with 95% confidence ellipses constructed around each treatment × day combination using a t-distribution. The percentage of fitted variance explained by each constrained axis was calculated from the eigenvalues of the constrained solution.

### Gabapentin treatment

Gabapentin (Product Number: G0318) was purchased from Tokyo Chemical Industry (TCI) and prepared in a 3mg/mL solution in 0.9% normal saline and dosed based off individual animal weights to achieve 30mg/kg administered intraperitoneally 45 minutes before behavioral recordings.

## Acknowledgements

The authors wish to thank Vivien Blecking and Anthony Menzel for their technical assistance. This work was supported by NIH Grants F31CA306387 (BARS), R37NS108278 (CLS), and R01NS070711 (CLS) and Advancing a Healthier Wisconsin (AHW) (5520857).

## Authorship contributions

BARS designed, performed, and analyzed experiments. BSD performed experiments. All authors interpreted the data and assisted with experiment design. BARS wrote the manuscript and CLS edited the manuscript. All figures were created by BARS. All experimental procedures, analysis, and manuscript drafts were supervised by CLS.

---

**Supplemental Figure 1.**
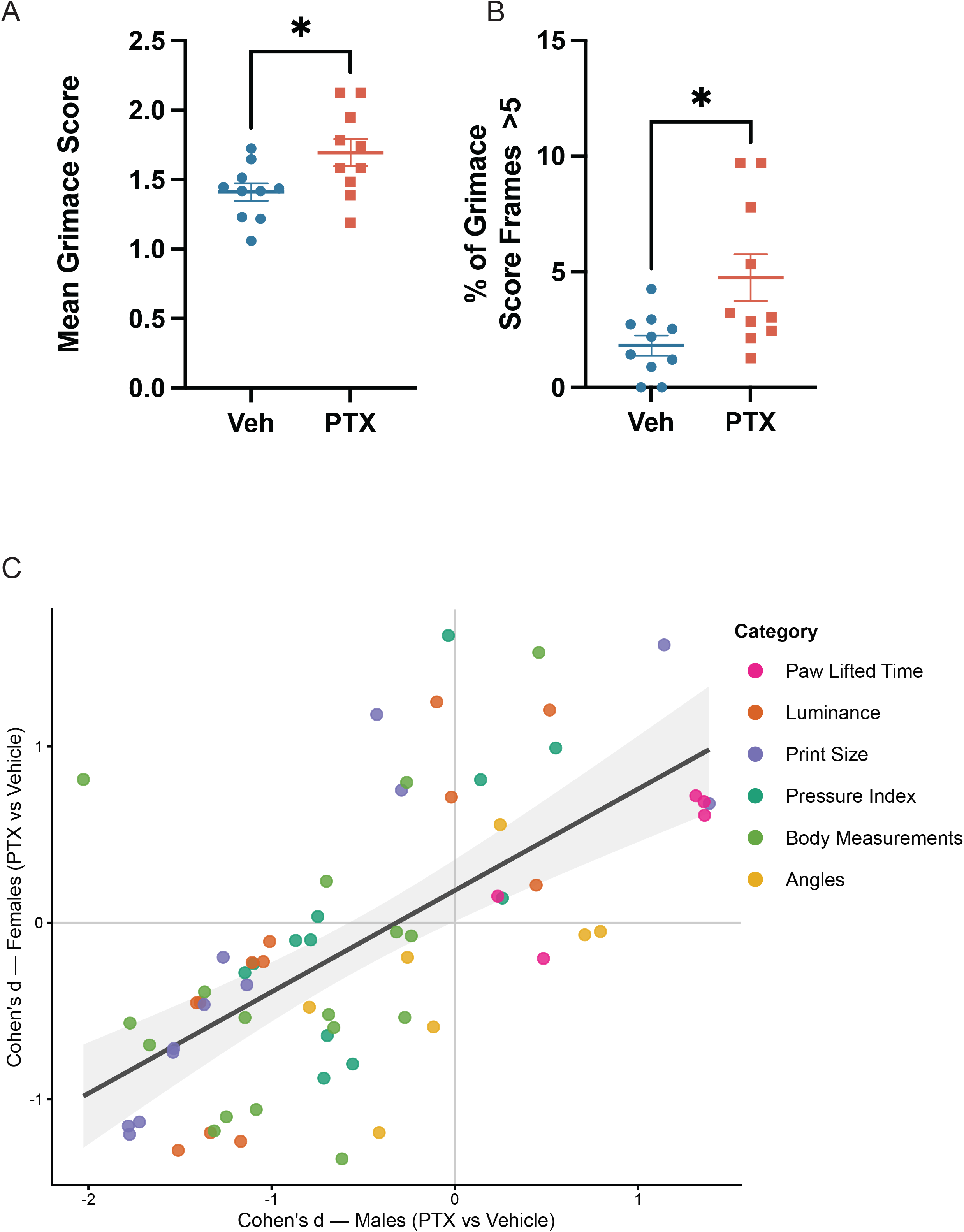
Conventional methods for detecting ongoing pain (facial grimace analysis), identify PTX animals during behavior recordings as having a higher mean grimace score (MGS) (A), and (B) spend more time in a high grimace state, consistent with an ongoing pain phenotype. C) Comparison of all analyzed 65 variables across sexes reveals the majority of variables have a consistent magnitude of effect in male and female PTX treated mice.

